# Convolutional Neural Network-Based Instance Segmentation Algorithm to Acquire Quantitative Criteria of Early Mouse Development

**DOI:** 10.1101/324186

**Authors:** Yuta Tokuoka, Takahiro G Yamada, Noriko F Hiroi, Tetsuya J Kobayashi, Kazuo Yamagata, Akira Funahashi

## Abstract

In embryology, image processing methods such as segmentation are applied to acquiring quantitative criteria from time-series three-dimensional microscopic images. When used to segment cells or intracellular organelles, several current deep learning techniques outperform traditional image processing algorithms. However, segmentation algorithms still have unsolved problems, especially in bioimage processing. The most critical issue is that the existing deep learning-based algorithms for bioimages can perform only semantic segmentation, which distinguishes whether a pixel is within an object (for example, nucleus) or not. In this study, we implemented a novel segmentation algorithm, based on deep learning, which segments each nucleus and adds different labels to the detected objects. This segmentation algorithm is called instance segmentation. Our instance segmentation algorithm, implemented as a neural network, which we named QCA Net, substantially outperformed 3D U-Net, which is the best semantic segmentation algorithm that uses deep learning. Using QCA Net, we quantified the nuclear number, volume, surface area, and center of gravity coordinates during the development of mouse embryos. In particular, QCA Net distinguished nuclei of embryonic cells from those of polar bodies formed in meiosis. We consider that QCA Net can greatly contribute to bioimage segmentation in embryology by generating quantitative criteria from segmented images.

## Introduction

The improved technologies for live cell imaging help to obtain multiple high-quality time-series three-dimensional (3D) fluorescence microscopic images [1–6]. In embryology, a number of studies have attempted to acquire quantitative criteria such as chromosome numbers, the synchrony of cell division, as well as determining the rate of development [7–9]. To analyze the time-series 3D microscopic images of developing embryos with fluorescently labeled nuclei, these studies used image segmentation. Segmentation methods in bioimage processing (such as filtering, thresholding, morphological operations, watershed transformation, and mask processing [4, 9–13]) require some parameter values. Because these methods are based on image-processing algorithms, they fail to detect an object in an image when this object does not fit the pattern described in the algorithm. Even though the optimal parameter values depend on the features of each image and the microscopy system, these values are arbitrarily set by the analyst, and further optimization tends to be neglected.

Previous studies have investigated embryonic development using different image processing methods [4, 9–13]. For time-lapse observation of early-stage *Drosophila* embryos, Keller et al. implemented digital scanned laser light sheet fluorescence microscopy with incoherent structured illumination microscopy (DSLM-SI) [4]. They performed nuclear segmentation of time-series images acquired by DSLM-SI. Their algorithm was based mainly on image processing. The images obtained by DSLM-SI have a high signal-to-noise ratio. *Drosophila* embryos are easily amenable to imaging because they are more transparent than embryos of other model organisms, such as mice. They were able to successfully perform segmentation of time-series images, using an imageprocessing algorithm, some limitations still remain. The segmentation accuracy dramatically decreased with embryo development: it was 95% at 2-4.5 h post-fertilization (h.p.f.), 73% at 4.5-7 h.p.f., and 54% at 7-11.5 h.p.f.. Analysis of time-series 3D fluorescence microscopic images is difficult because (1) fluorescence intensity decreases along the *z*-axis because the inner part of the embryo is not transparent: (2) fluorescence intensity decreases with time because of the fading of the fluorophore: and (3) high spatial resolution cannot be achieved because a balance between cytotoxicity and photographing speed needs to be maintained. The current low accuracy can be attributed to the fact that the change in the spatiotemporal features of time-series 3D fluorescence microscopic images is not correctly grasped.

Convolutional Neural Network (CNN), a deep learning algorithm, has been proposed [14–20]. In image analysis, CNN performs better than other methods [21]. A critical advantage of CNN is automatic extraction of image features useful for analysis. CNN has also been applied to bioimage segmentation methods, and its performance was superior to that of the previous methods [14–18, 20]. Çiçek et al. implemented 3D U-Net based on CNN and used it for segmentation of microscopic images of *Xenopus* kidney tissue [15]. The authors produced training data by manually annotating each image voxel-wise with “kidney tubule”, “inside kidney tubule”, or “background”. As a result of learning these training data, a high value of Intersection over Union (IoU), an evaluation metric for segmentation, was achieved (0.723), indicating high accuracy of 3D U-Net. Ho et al. developed a 3D CNN algorithm and used it to perform segmentation of 3D fluorescence microscopic images of labeled nuclei of rat kidney by the algorithm [18]; the algorithm achieved a voxel accuracy of 0.922. However, the remaining problem of the above algorithms is that some segmented nuclear regions are fused with other regions, which disturbs acquisition of quantitative criteria from bioimages.

The above segmentation algorithms are based on Fully Convolutional Networks (FCN) [22], which consist only of convolution layers in CNN, and the segmentation methodology of FCN is called semantic segmentation. Because semantic segmentation assigns the same label to the objects of the same class (Fig. 1), the regions are fused when neighboring or overlapping objects are segmented [23]. Semantic segmentation is appropriate for objects at the tissue level, but not for objects at the cellular or organelle level. In this study, we focused on the other segmentation methodology, called instance segmentation [24], adds a different label to each object of the same class (Fig. 1) and is suitable for segmentation of cells and nuclei. Because individual objects have to be recognized, instance segmentation is a more difficult task than semantic segmentation. Many instance segmentation methods using CNN proposed in the field of general image recognition are complicated and difficult to apply to other fields [24–26]. Almost all of these methods target 2D images, and no practical algorithm is able to execute instance segmentation for 3D bioimages.

**Figure 1:**
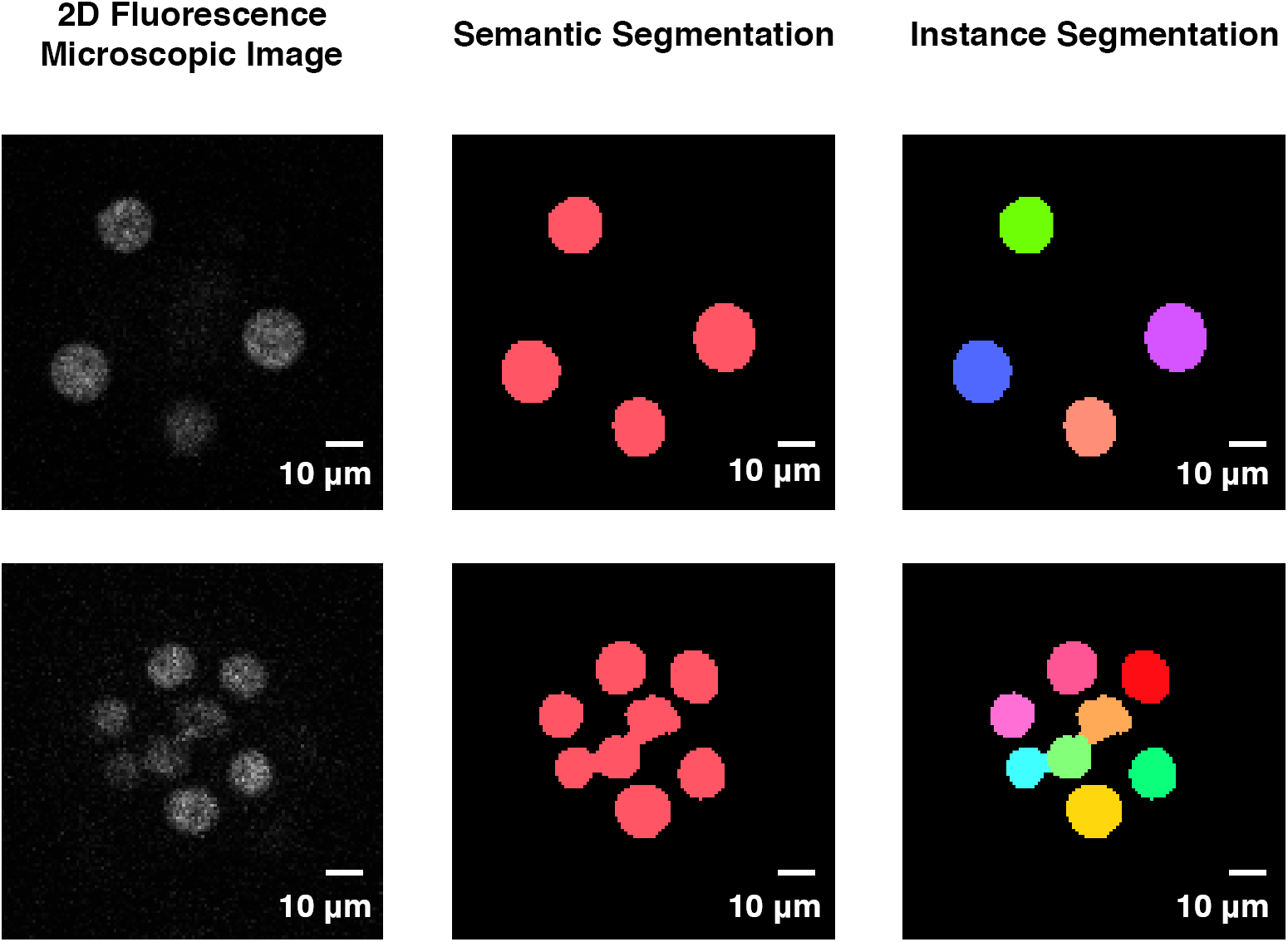
Conceptual diagram of different segmentation methods. In a 2D fluorescence microscopic image, all objects to be segmented are of the same class (nucleus). Semantic segmentation assigns the same label to all objects of the same class, while instance segmentation assigns different labels. When the objects are sufficiently separated in space (upper panels), segmentation of both types is accurate. When objects are adjacent or overlap (lower panels), semantic segmentation fuses the object regions, but instance segmentation does not.

Here, we developed QCA Net, a new CNN-based segmentation algorithm, which does not cause object fusion. QCA Net has a simple structure, combining conventional semantic segmentation algorithms, and can be easily applied to bioimage analysis. We trained QCA Net using part of a single early-stage mouse embryo and performed instance segmentation of 11 mouse embryo images. We accurately acquired the shape of the nucleus without nuclear region fusion and extracted quantitative criteria (timeseries data for nuclear number, volume, surface area, and center of gravity coordinates). QCA Net not only performed nuclear segmentation accurately but also excluded the nuclei of polar bodies, which are difficult to distinguish in image processing.

## Results

### Quantitative Criterion Acquisition Network (QCA Net)

We called the implemented algorithm Quantitative Criterion Acquisition Network (QCA Net); it performs instance segmentation of 3D fluorescence microscopic images (Fig. 2). QCA Net consists of two subnetworks: Nuclear Segmentation Network (NSN) and Nuclear Detection Network (NDN). NSN learns the task of nuclear segmentation, and NDN learns the task of nuclear identification. The inputs of QCA Net are time-series 3D fluorescence microscopic images. QCA Net executes instance segmentation of the images at each time point. An example of instance segmentation at the 4-cell stage is shown in Figure 2 (lower panel).

**Figure 2:**
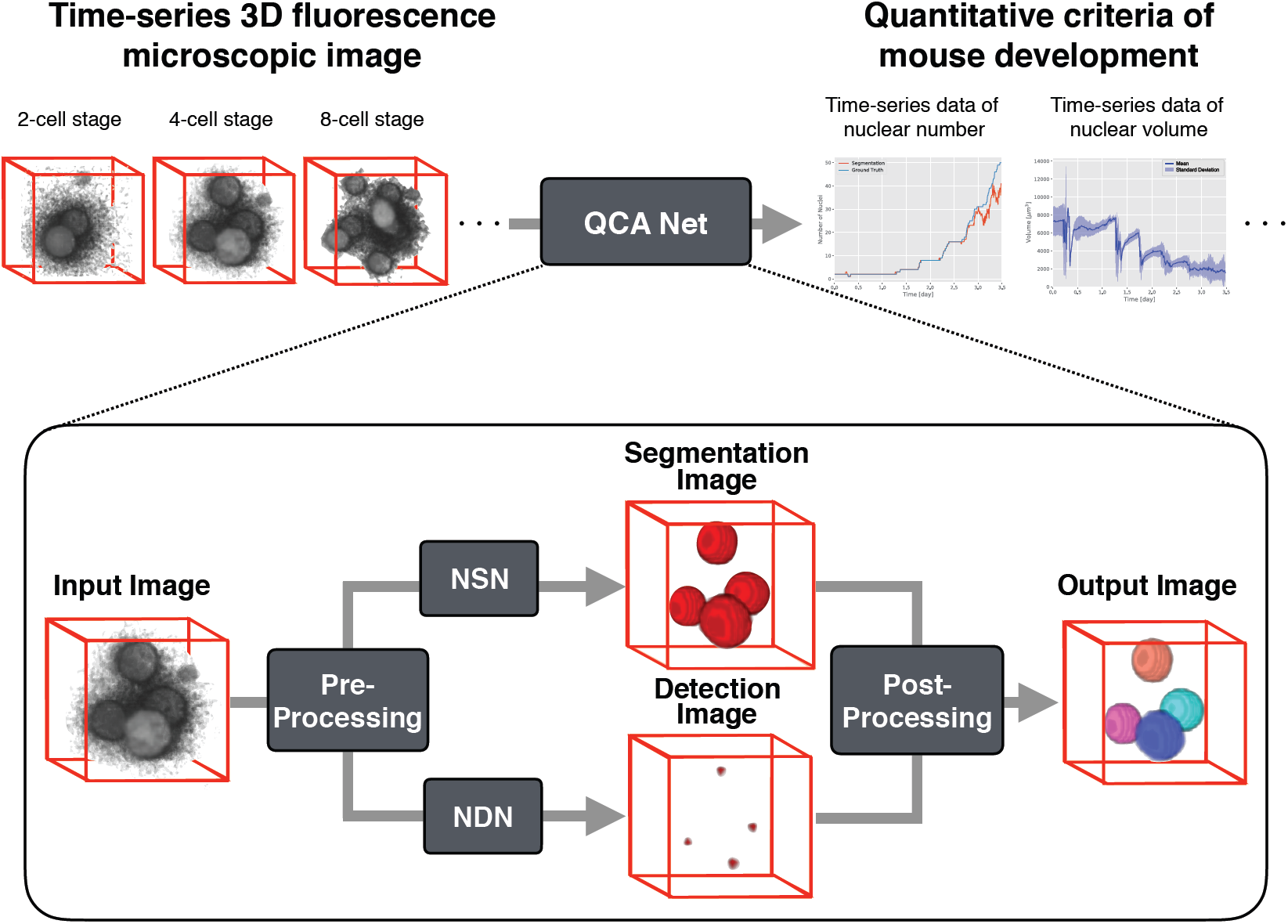
Flow diagram of our implementation of the QCA Net algorithm. QCA Net extracts quantitative criteria of mouse development from time-series 3D fluorescence microscopic images of earlystage mouse embryos as input. QCA Net first performs pre-processing of the input image at each time point. The pre-processed image is then processed in parallel by the Nuclear Segmentation Network (NSN), which segments the nuclei, and the Nuclear Detection Network (NDN), which identifies them. The nuclear region segmented by NSN is divided by marker-based watershed in post-processing using the nuclear center identified by NDN. By performing instance segmentation at each time point, QCA Net acquires a time-series instance segmentation image, which is used to extract the quantitative criteria of mouse development.

To achieve instance segmentation at each time point, the following procedure is used. Initially, the input image is pre-processed. Next, NSN performs semantic segmentation of the nuclear region, and NDN performs semantic segmentation of the center of this region in parallel. Finally, the nuclear region estimated by NSN is segmented by marker-based watershed from the center identified by NDN.

Sequential input of time-series images to QCA Net outputs instance segmentation images at each time point (Fig. 2, upper panel). QCA Net can extract the time-series data for the nuclear number, volume, surface area, and center of gravity coordinates. QCA Net is open source software and can be downloaded from https://github.com/funalab/QCANet.

To qualitatively evaluate the performance of QCA Net, we compared its segmentation accuracy with that of 3D U-Net [15], which is a semantic segmentation algorithm for 3D bioimages. QCA Net and 3D U-Net learned the same dataset, which was sampled from a single early-stage mouse embryo, whose cell nuclei were fluorescently labeled with mRFP1 fused to histone 2B, which is a chromatin marker [1]. QCA Net and 3D U-Net then performed segmentation of the embryo that was used for learning but at time points different from the embryo used for learning. The 3D fluorescence microscopic images of the segmentation target were those of a mouse embryo at the 16-cell and 50-cell stages whose cell nuclei were fluorescently labeled (Fig. 3a,d). QCA Net determined the nuclear regions correctly and accurately performed instance segmentation (Fig. 3b,e), whereas the segmented nuclear regions determined by 3D U-Net were fused, indicating that 3D U-Net did not perform segmentation accurately.

**Figure 3:**
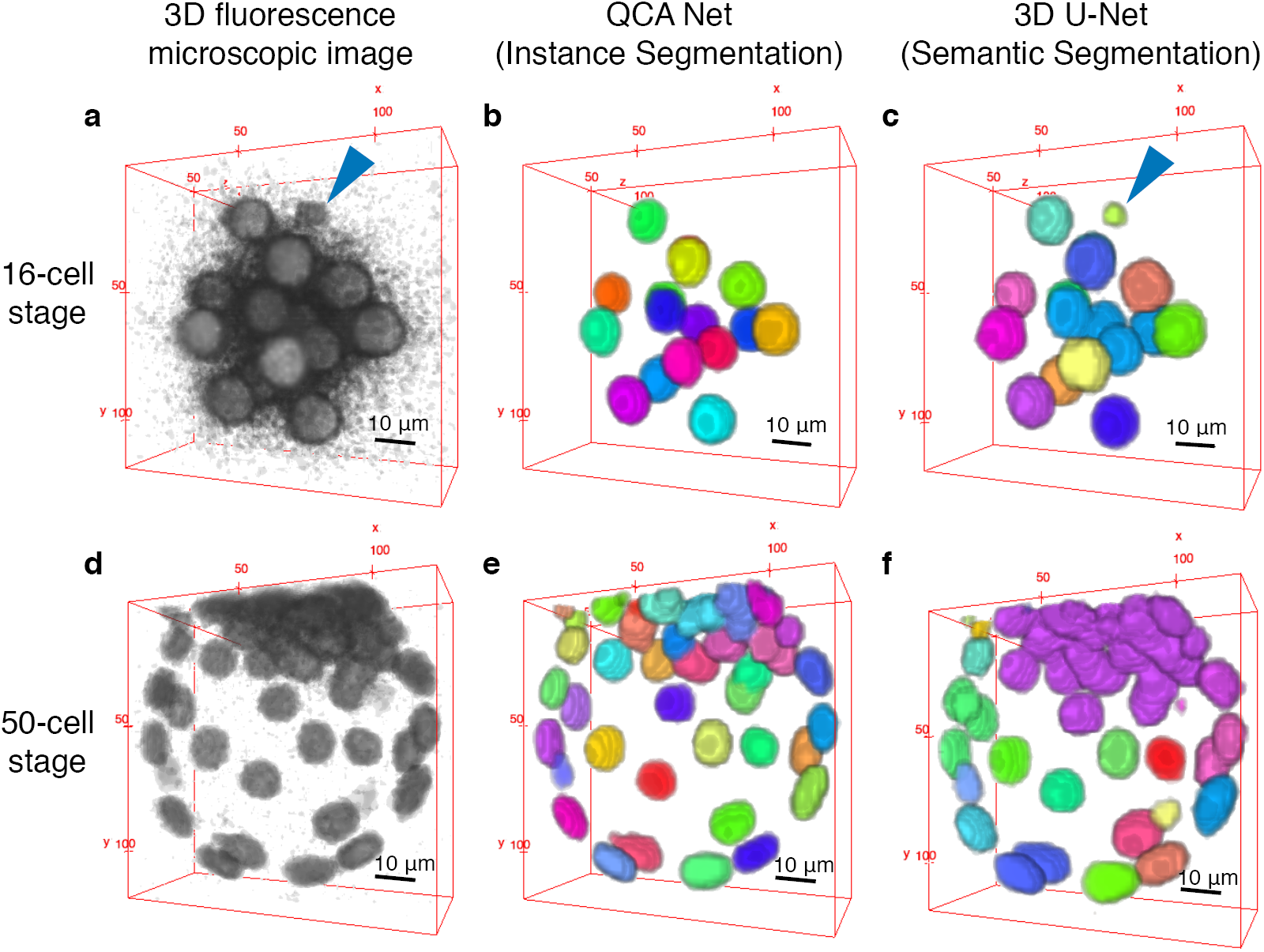
Comparison of segmentation by QCA Net and 3D U-Net. Segmentation of 3D fluorescence microscopic images of one mouse embryo at two cell stages is shown. Each color represents an individual segmented nuclear region. Images of a mouse embryo about 2.4 days (16-cell stage) and about 3.5 days (50-cell stage) after the pronuclear stage were used. QCA Net performed accurate segmentation without nuclear region fusion and excluded nuclei of a polar body (blue arrowhead), whereas the fusion of nuclear regions and segmentation of the nuclei of the polar body occurred in 3D U-Net.

Because polar bodies have nuclei, they are fluorescently labeled, and it is difficult to exclude them from segmentation results by using traditional image processing algorithms. Aiming at acquiring the quantitative criteria of mouse development, we did not add the nuclei of polar bodies to training data as the ground truth and confirmed that the nuclei of polar bodies were excluded from segmentation (Fig. 3b). Therefore, QCA Net can recognize and distinguish the nuclei of embryonic cells and those of polar bodies. Although 3D U-Net learned the same training data as QCA Net, 3D U-Net failed to exclude the nuclei of polar bodies from segmentation (Fig. 3c).

QCA Net accurately performed segmentation of time-series images of the mouse embryo used for learning (Supplementary Video 1). This was particularly challenging because early-stage mouse embryos form blastocysts whose cells, and therefore nuclei, are located very close to each other.

QCA Net and 3D U-Net also performed segmentation of 10 different mouse embryos, which were not included in the learning datasets, at the 8-, 16-, and 32-cell stages; cell nuclei were fluorescently labeled (Supplementary Fig. 1a,d,g,j). Despite the differences between these embryos and the embryo used for learning, QCA Net accurately performed segmentation of the nuclear regions (Supplementary Fig. 1b,e,h,k) and also excluded the nuclei of polar bodies (blue arrowheads in Supplementary Fig. 1a,d,g). However, the segmentation by 3D U-Net had marked fusion of the nuclear regions (Supplementary Fig. 1c,f,i,l), and the nuclei of polar bodies were also segmented as nuclei of cells (blue arrowheads in Supplementary Fig. 1c,f,i). The segmented nuclei of polar bodies were one of the causes for the false-positive errors. Supplementary Video 2 shows time-series segmentation results of 10 different mouse embryos.

Developing mouse embryos have complex characteristics, such as the rate of development, nucleus arrangement, shape, and fluorescence intensity, but QCA Net robustly performed instance segmentation.

### Quantitative evaluation of QCA Net

We quantitatively compared the segmentation accuracy of QCA Net and 3D U-Net. An answer was considered correct when a voxel of an object region was classified as such (true positive, TP) or a voxel of a background region was classified as such (true negative, TN). An answer was considered incorrect when a voxel of a background region was classified as an object region (false positive, FP) or a voxel of an object region was classified as a background region (false negative, FN). Accordingly, the evaluation metric of segmentation was defined as 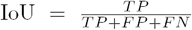, where TP, FP, and FN denote the numbers of voxels defined above. IoU is conventionally used in segmentation because it measures the false-positive and false-negative rates comprehensively [15]. However, because IoU was calculated for each image, it could not evaluate whether (or not) segmentation was accurate without fused nuclei and was thus not suitable for evaluating instance segmentation. Therefore, we also used a metric called MUCov [24], which is based on IoU, for the evaluation of instance segmentation. 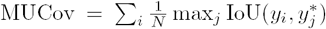, where *N* is the number of segmented nuclei, *y* is the segmented nuclear region, *y*^*∗*^ is the ground truth of the nuclear region, *i* is the label attached to the segmented nuclear region (*i* = 1,*…, N*), and *j* is the label attached to the ground truth of the nuclear region.

To evaluate both semantic segmentation (3D U-Net and QCA Net without NDN, which includes preprocessing, NSN, and post-processing) and instance segmentation (QCA Net), we used IoU and MUCov. In these evaluations, we used images of the mouse embryo used for learning taken at 18 time points of (Table 1). The average value of each metric at each time point and its standard deviation are shown in Table 1.

**Table 1:** Quantitative evaluation of segmentation accuracy. The evaluated objects were embryos imaged at 18 time points; the same images were also used as the ground truth for learning. IoU was used as a metric for semantic segmentation, and MUCov was used as a metric for instance segmentation. The values are mean and standard deviation.

| Model | IoU | MUCov |
| --- | --- | --- |
| QCA Net (instance segmentation) | 0.817 (0.121) | 0.801 (0.117) |
| QCA Net without NDN (semantic segmentation) | 0.813 (0.121) | 0.647 (0.130) |
| 3D U-Net (semantic segmentation) | 0.665 (0.123) | 0.334 (0.120) |

The IoU value of QCA Net exceeded that of 3D U-Net, which has been reported to accurately perform semantic segmentation [15]. Therefore, QCA Net was superior to the currently existing algorithm of segmentation, likely because QCA Net performed parameter tuning using Bayesian optimization [27] and because hyperparameters of QCA Net used for learning (e.g., optimization and regularization) differed from those of 3D U-Net. The IoU value of QCA Net slightly exceeded that of QCA Net without NDN because QCA Net succeeded in exclusion of false-positive nuclei using identification by NDN.

The MUCov value of QCA Net exceeded those of both QCA Net without NDN and 3D U-Net. The MUCov values of QCA Net without NDN and 3D U-Net were likely reduced by the fusion of nuclear regions in the segmentation (Fig. 3 and Supplementary Fig. 1). NDN identified nuclei with high accuracy, which might have enabled QCA Net to precisely divide the fused nuclear regions. We evaluated the nuclear identification accuracy of NDN using F-measure (Supplementary Note 1 and Supplementary Table 1). F-measure comprehensively measures the false-positive and false-negative rates to evaluate nuclear identification. The average F-measure of 11 different mouse embryos was 0.915 in NDN, higher than that (0.856) obtained with a previous algorithm [9] for our datasets; although the algorithm used in the previous study identified nuclei with high accuracy, the accuracy of NDN was superior.

These results quantitatively show that QCA Net performed instance segmentation with high accuracy. Evaluation at each time point is shown in Supplementary Tables 2 and 3.

### Acquisition of time-series data of quantitative criteria of mouse development by QCA Net

We extracted the data of the critical quantitative criteria used in standard embryonic development analysis: the nuclear number, volume, surface area, and center of gravity coordinates. The data of the nuclear number are shown in Figure 4a. Each series represented the result by QCA Net and the ground truth. The F-measure of nuclear identification accuracy (0.932) exceeded 0.89810. Therefore, QCA Net extracted the time-series data of nuclear number with high accuracy.

**Figure 4:**
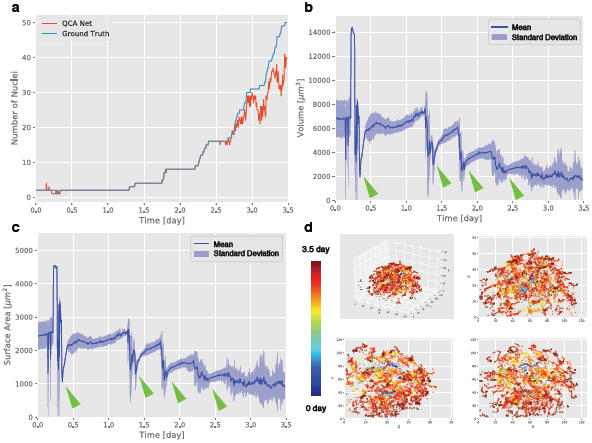
Quantitative criteria of mouse development extracted by QCA Net from time-series data. (a) Nuclear number. (b) Nuclear volume. The nuclear volume tended to rapidly decrease and then recover (green arrowheads). This tendency may indicate the anaphase. (c) Nuclear surface area. Similar to (b), this time course also captures the features of anaphase (green arrowheads). (d) The nuclear center of gravity coordinates. The upper left panel shows the results displayed in 3D; upper right, *XZ* cross-section; lower left, *YZ* cross-section; lower right, *XY* cross-section. Color shift from cold to warm indicates the course of development. With time, the internal clearance is widening, indicating blastocyst formation.

The data of the nuclear volume are shown in Figure 4b. We found a periodical tendency of sharp decreases in nuclear volume followed by its recovery. In *Drosophila* embryos [28], a similar tendency was caused by chromosome condensation in anaphase. Thus, QCA Net appeared to correctly extract a specific characteristic of anaphase. The nuclear volume from the pronuclear to 2-cell stage (0-1.3 days) was about 7,000 *μ*m^3^ (Fig. 4b). The volume of the mouse embryo at the 2-cell stage has been determined as approximately 56,000 *μ*m^3^ [29]. Therefore, the scale of the nucleus volume was within that of the mouse embryo, and our value of the nuclear volume was reasonable. The accuracy of QCA Net for nuclear volume is mostly affected by the accuracy of instance segmentation, which is evaluated by MUCov. The MUCov value of QCA Net (0.801) shows that QCA Net accurately extracted the time-series data of nuclear volume.

The data of the nuclear surface area are shown in Figure 4c. The tendency was similar to that described above for the nuclear volume, probably because the nucleus is spherical. Thus, QCA Net accurately extracted the time-series data of the nuclear surface area.

The data of the nuclear center of gravity coordinates are shown in Figure 4d. During the development from morula to blastocyst, the internal space expands. Cells of the outer layer of the blastocyst (trophectoderm [30]) becomes the source of extraembryonic tissue. We observed an expansion of the internal space during blastocyst formation. The time-series data of the nuclear center of gravity coordinates accurately extracted by QCA Net showed this phenomenon.

Quantitative criteria of 10 different mouse embryos that were not used in learning are shown in Supplementary Figs. 2-5). The results were similar to those of evaluation for the images used for learning and indicated that QCA Net was able to acquire quantitative criteria from mouse embryos that were not used for learning.

### Recognition of polar bodies

In early-stage embryos, polar bodies are formed in oocyte meiosis. Polar bodies have a nucleus but hardly any cytoplasm, and they slowly degenerate during development and disappear naturally. Therefore, polar bodies may not be related to normal development and should be excluded from segmentation targets. However, polar bodies tend to be extracted by image processing because microinjection of mRNA encoding a fluorescent protein results in its production in polar bodies.

QCA Net segmented nuclei except those of polar bodies (Fig. 5). In the first case, NSN and NDN excluded the nuclei of polar bodies (Fig. 5a-c). In the second case, NSN identified the nuclei of polar bodies, but NDN excluded them (Fig. 5d-f). Despite this false-positive error of NSN, the segmentation region of the nuclei of polar bodies was excluded in post-processing owing to their exclusion by NDN.

**Figure 5:**
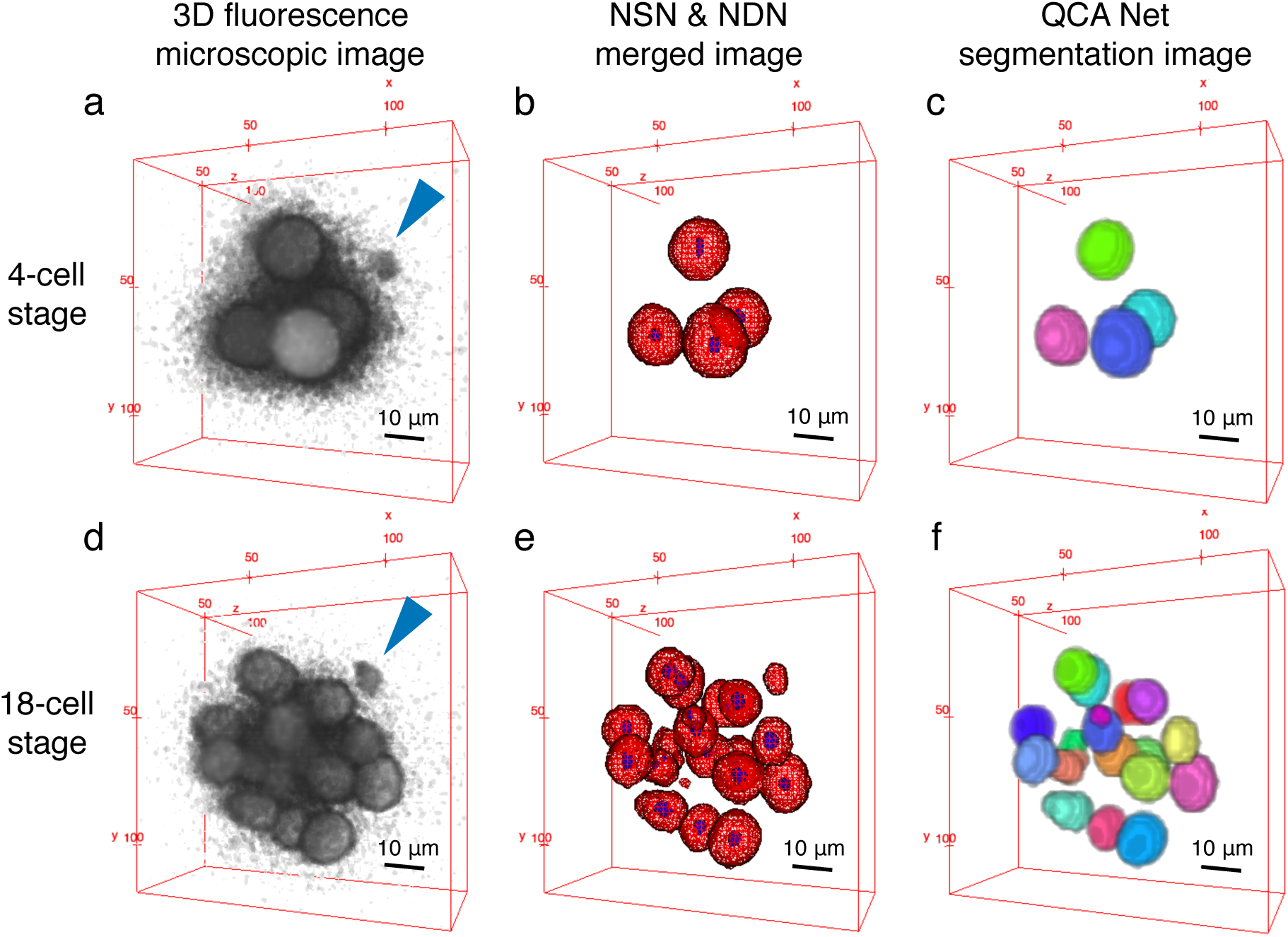
Polar bodies are recognized and excluded from segmentation targets. (a-c) Both NSN and NDN excluded the polar body (blue arrowhead). (d-f) NSN performed segmentation of the polar body, but NDN excluded it. The 3D images were acquired at (a) the 4-cell stage and (d) 18-cell stage. (b, e) Nuclear segmentation by NSN (red mesh) and nuclear identification by NDN (blue mesh). (c, f) Instance segmentation by QCA Net excluded polar bodies in both cases.

We considered why the high accuracy of segmentation in QCA Net was achieved in the second case. QCA Net independently identified nuclei by NSN and NDN. The watershed process using the result of NDN was performed to divide the nuclear region segmented by NSN. Therefore, when NDN identified the false-positive error of NSN, the nuclei of polar bodies were excluded in post-processing. These results show that QCA Net performs high-quality analysis of bioimages.

## Discussion

Segmentation is an important and challenging task of bioimage analysis aimed at uncovering biological phenomena, such as embryogenesis. Although segmentation accuracy has been improved by methods based on the deep learning algorithms [14–18], these methods are based on semantic segmentation and many problems remain, for example fusion of objects. In this study, we developed QCA Net, which is an instance segmentation algorithm for high-density cells or intracellular organelles in 3D images. QCA Net has a simple structure so that the algorithm can be readily applied to various microscopic images such as bright field, phase contrast, and differential interference contrast. In particular, NDN could estimate the central nuclear region because cells and intracellular organelles have a characteristic patterns and shapes. Because the NDN subnetwork is simple and powerful in bioimage analysis, there is no need to use a complicated object detection algorithm, such as those used in the field of general image recognition [31, 32].

In comparison with semantic segmentation, QCA Net was especially effective, both qualitatively and quantitatively, for high-density objects. The segmentation accuracy of QCA Net (IoU, 0.817; MUCov, 0.801) exceeded that of the semantic segmentation algorithm 3D U-Net [15] (IoU, 0.665; MUCov, 0.334). The nuclear identification accuracy of NDN (F-measure, 0.915) exceeded that in the previous study [9] (F-measure, 0.856). Using the segmentation images, we extracted the quantitative criteria of mouse development such as nuclear number, volume, surface area, and center of gravity coordinates were extracted. In development, cell division patterns follow specific rules, although some fluctuations of this pattern are allowed. Information on the quantitative criteria will be essential for uncovering the robust mechanisms of development. We expect that QCA Net will considerably improve the quality and throughput of analysis in embryology.

The role of polar bodies in the development have been discussed for a long time [33, 34], but there is no clear answer as to why polar bodies exist in the embryo. Because QCA Net recognized the nuclei of polar bodies, it seems possible to trace only polar bodies during development; thus, QCA Net will be a powerful tool in developmental biology. Yet, the way how QCA Net recognizes the polar bodies was not evident because the regression of deep learning was too complicated. Some studies have tried to analyze learned features [35, 36]. It was reported that each layer in the neural network has a role in image-processing such as filtering [37]. Based on these studies, the regression by deep learning could be replaced by a combination of image-processing. If this combination is revealed and the layer which has a role in distinguishing nuclei of embryonic cells and those of polar bodies is determined, the way of recognition of polar bodies will be uncovered.

## Methods

### Fluorescence imaging for learning and evaluation

The 5,522 time-series images of 11 early mouse embryos from the pronuclear stage to the maximum of the 53-cell stage were taken under a 3D confocal fluorescence microscope. The conditions of image acquisition are shown in Supplementary Table 4. Each embryo had a different developmental rate and was at a different developmental stage (Supplementary Fig. 6).

### Ground truth creation

We manually created the ground truth from fluorescence microscopic images at 18 time points in a single embryo using Fiji [38]. NSN and NDN learned the tasks of nuclear segmentation and detection, respectively from the created ground truth. This embryo developed from the pronuclear stage to the 50-cell stage. We excluded the nuclei of polar bodies from the ground truth. The ground truth to learn the task of nuclear identification was a spherical region with a diameter of 5 voxels; this region was created based on the nuclear center of gravity coordinates. The sampled 18 time points and the developmental stages are shown in Supplementary Table 5. The ground truth, which compared the time-series data of the nuclear number, was created at each time point for 11 mouse embryos as shown in Supplementary Figure 7.

### QCA Net overview

QCA Net consists of NSN, which learned the nuclear segmentation task, and NDN, which learned the nuclear identification task (Fig. 2). QCA Net performs instance segmentation of the time-series 3D fluorescence microscopic images at each time point, and the quantitative criteria for mouse development are extracted from the acquired time-series segmentation image.

We implemented QCA Net in Python 2.7 and used Chainer [39], which is an open-source deep learning framework. We used NVIDIA Tesla K40 (operating frequency, 745 MHz; single precision floating point performance, 4.29 TFLOPS) and NVIDIA Tesla P100 (1189 MHz, 9.3 TFLOPS) for calculation of learning and segmentation. P100 is on Reedbush-H, a calculation server of the University of Tokyo Information Infrastructure Center. The source code of QCA Net is available from https://github.com/funalab/QCANet.

### Pre-processing in QCA Net

The objective of normalization was to prevent divergence of values and gradient disappearance in learning. The value of each voxel to be normalized (*I*′) is defined by

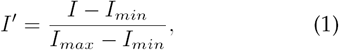

where *I* is the value of each voxel to be normalized, *I*_*max*_ is the maximum voxel value in the image, and *I*_*min*_ is the minimum voxel value in the image. The value of *I*′ is obtained for all the voxels in the image, and the range of the voxel values is [0, 1].

Mirror padding is a method for padding around the image. The objective of mirror padding was to fit the patch area within the image even if the voxel of interest is out of the image. Mirror padding was performed by acquiring voxel values inside from *m* pixels from the edge of the image and extrapolating this mirror image to the outer edge. The patch size of QCA Net was 128 voxels, so the size of mirror padding was 64 voxels.

Since *x, y*, and *z*-axis resolution in the microscopic image to be analyzed was 0.8 : 0.8 : 1.75 *μ*m (Supplementary Table 4), it was necessary to change it to the actual scale ratio of 1 : 1 : 1. We interpolated 2.1875 times in the *z*-axis direction as bicubic interpolation.

In general, when the number of training data samples is small, a method of processing data and increasing the number of samples to prevent overlearning is used. In this study, the number of samples of the ground truth was small, so we performed data augmentation and increased the number of data four times for each training image by flipping on the *x*-axis, the *y*-axis, and both the *x* and *y*-axes. Since the bias of the luminance in the *z*-axis direction, which is a feature of a time-series 3D fluorescence microscopic image, is always constant, we did not expand the data in this direction.

### Nuclear Segmentation Network

NSN performs semantic segmentation of nuclear regions from 3D fluorescence microscopic images. We used Stochastic Gradient Descent (SGD) as an optimization method for learning. The structure of the network is based on 3D U-Net [15], and parameter tuning suitable for the dataset was performed by the Bayesian optimization in SigOpt (https://sigopt.com). NSN had 1,146,896 parameters fewer than in 3D U-Net (Supplementary Table 6).

### Nuclear Detection Network

NDN is a network that performs semantic segmentation of nuclear center regions from 3D fluorescence microscope images. We used Adam [40] as an optimization method for learning. The structure of the network was based on 3D U-Net [15], and parameter tuning suitable for the dataset was performed by the Bayesian optimization in SigOpt. NDN had 44,447,940 parameters more than in 3D U-Net (Supplementary Table 7).

### Post-processing in QCA Net

We performed (a) reinterpolation and (b) marker-based watershed on the semantic segmentation image output from NSN and NDN. Reinterpolation restores the resolution of the image interpolated for segmentation and identification. Marker-based watershed divides the semantic segmentation region by watershed with the center region of the identified nucleus as a marker. Post-processing enabled QCA Net to execute instance segmentation.

### Tuning the hyperparameters of NSN & NDN

Hyperparameters in NSN and NDN (10 in total) were optimized by the Bayesian optimization (Supplementary Table 8). SigOpt was used as the optimization platform.

The following hyperparameters were optimized. Up- and downsampling denotes the number of times to perform upsampling and downsampling. Initial channels denotes the number of channels in the first convolutional layer. The number of channels in the subsequent convolutional layers was based on the structure of 3D U-Net, which is doubled in the upsampling section and is halved in the downsampling section. Kernel size denotes the common kernel size in all convolutional layers. Weight decay denotes the parameter of the L2 norm.

In this experiment, we used SGD and Adam as the optimizers. The initial learning rate, which is a parameter of SGD, denotes the initial value of the learning coefficient, and the decay learning rate denotes a parameter for multiplying the initial learning rate when the accuracy of the reference evaluation metrics is lowered. The reference evaluation metrics were IoU in NSN and F-measure in NDN. Adam’s, *α, β*_1_, *β*_2_ and *ϵ* are parameters incorporated into the expression that determines the learning rate of Adam. Epoch denotes the number of learning iterations; an attempt to input all the training data and make neural network learn is defined as one epoch. Epoch was fixed at 50 for optimization. Since Epoch directly affects the amount of computation, we consider that if computation resources are abundant, it should be added to the optimization target.

### Model architecture and learning conditions of NSN

NSN hyperparameters were determined by the Bayesian optimization (Supplementary Tables 9 and 10), and the model architecture of NSN was determined based on these hyperparameters (Supplementary Table 6). The output function of NSN, called softmax, is defined by

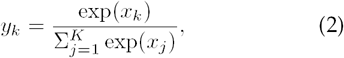

where *K* denotes the number of classes (nucleus or background region), *x* denotes each input from the final layer, and *y* denotes the output value. The objective function of NSN, softmax cross entropy, is defined by

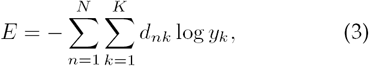

where *d* denotes the ground truth and *N* denotes the number of learning data. Epoch was fixed at 150 for learning. Using SGD and Adam, we evaluated learning the model (Supplementary Fig. 8). Because NSN trained by SGD performed nuclear segmentation with high accuracy, we adopted SGD-trained NSN for QCA Net.

### Model architecture and learning conditions of NDN

NDN hyperparameters were determined by the Bayesian optimization (Supplementary Tables 11 and 12), and the model architecture of NDN was determined based on these hyperparameters (Supplementary Table 7). Similar to NSN, softmax and softmax cross entropy were used as the output and objective functions, respectively. Epoch was fixed at 150 for learning. Using SGD and Adam, we evaluated learning the model by NDN (Supplementary Fig. 9). Because NDN trained by Adam performed nuclear identification with high accuracy, we adopted Adam-trained NDN for QCA Net.

### Extraction of quantitative criteria from segmentation image

Nuclear number was extracted by counting the number of labels in segmentation images. Nuclear volume was extracted by converting the voxel number of the segmented nuclear region for each label to the actual scale. Nuclear surface area was extracted by converting the voxel number of the nuclear region that was in contact with the background region to the actual scale. The nuclear center of gravity coordinates were calculated as the center of gravity of the segmented nuclear region for each label.

## Acknowledgements

The author would like to thank NM Drissi for constructive criticism of the manuscript. The research was funded by a JSPS KAKENHI Grant (Number 16H04731). Computations were primarily performed using the computer facilities at The University of Tokyo (Reedbush). The Bayesian optimization was performed using SigOpt.

## Author contributions

Y.T. and A.F. designed the conceptual idea and the study. Y.T. implemented the algorithm of QCA Net. T.J.K. provided the ground truth for nuclear identification. K.Y. provided the datasets of mouse embryos. Y.T., T.G.Y., N.F.H., and A.F. wrote the manuscript with suggestions from the other authors.

